# A Practical Approach for Targeting Structural Variants Genome-wide in Plasma Cell-free DNA

**DOI:** 10.1101/2023.10.25.564058

**Authors:** Michael M. Murata, Fumie Igari, Ryan Urbanowicz, Lila Mouakkad, Sungjin Kim, Zijing Chen, Dolores DiVizio, Edwin M. Posadas, Armando E. Giuliano, Hisashi Tanaka

**Affiliations:** Department of Surgery, Cedars-Sinai Medical Center, West Hollywood, CA 90048, USA; Cedars-Sinai Cancer, Cedars-Sinai Medical Center, West Hollywood, CA 90048, USA; Department of Breast Oncology, Juntendo University, Tokyo, Japan; Department of Computational Biomedicine, Cedars-Sinai Medical Center, West Hollywood, CA 90048, USA; Department of Medicine, Cedars-Sinai Medical Center, West Hollywood, CA 90048, USA; Department of Biomedical Sciences, Cedars-Sinai Medical Center, West Hollywood, CA 90048, USA

**Author notes:** Corresponding author: Hisashi Tanaka. These authors contributed equally to this work (Michael M. Murata, Fumie Igari).

## Abstract

Interrogating gene mutations and aberrant DNA methylation in plasma cell-free DNA (cfDNA) has become increasingly common for monitoring tumor burden in cancer patients. However, no tests currently target chromosomal structural variants (SVs) genome-wide. Here, we report a simple molecular and sequencing workflow, Genome-wide Analysis of Palindrome Formation (GAPF-seq), to probe DNA palindromes, a type of SV that often demarcates gene amplification. Low-coverage next-generation sequencing of palindrome-enriched DNA uncovered skewed chromosomal distributions of high-coverage 1-kb bins (HCBs) in tumor DNA. When combined with traditional machine learning, GAPF-seq differentiated 39 breast tumors from matched control DNA with an Area Under the Curve (AUC) of 0.955. A proof-of-concept liquid biopsy study using cfDNA from 27 prostate cancer patients and 24 control individuals yielded an average AUC of 0.896. HCBs on the X chromosome emerged as a nearly decisive feature and were linked to androgen receptor gene amplification. As a simple and agnostic liquid biopsy approach, GAPF-seq could fill this technological gap, offering unique cancer-specific SV profiles.

## Introduction

Genome instability is a defining hallmark of cancer.^1–5^ Changes at the single nucleotide level (i.e., mutations) can activate oncogenes or inactivate tumor suppressor genes. Equally prevalent are structural rearrangements of large chromosomal segments, referred to as structural variants (SVs), such as duplications, deletions, and translocations. These genetic changes are the primary drivers of malignant transformation, disease progression, and therapy resistance. Beyond these roles in cancer biology, tumor-specific DNA changes can serve as biomarkers for cancer detection, a field that has gained momentum with liquid biopsy.^6,7^ Liquid biopsy holds promise as a less invasive and more repeatable alternative to currently available methods in the clinic, such as needle biopsies or imaging scans. Traditionally, blood has been a source of protein biomarkers such as prostate-specific antigen (PSA).^8^ More recently, with the advent of DNA analysis technologies for cancer genomes, DNA released into the blood from tumor cells, termed circulating tumor DNA (ctDNA), has become a mainstream target for developing liquid biopsy-based clinical tests. Several Food and Drug Administration (FDA) approved tests are currently in clinical use.^9–14^ These tests identify tumor-specific mutations in cell-free DNA (cfDNA) isolated from plasma, either using a single-gene platform or Next-Generation Sequencing (NGS) to target multiple cancer genes (i.e., targeted sequencing). While reporting residual tumor burden and actionable mutations for advanced cancer patients, there is potential for extend mutation detection in plasma cfDNA to cancer screening in the general population. However, even non-malignant hematopoietic cells accumulate clonal mutations in cancer-associated genes such as *p53* and *KRAS*.^15–17^ This phenomenon, termed clonal hematopoiesis, complicates the use of mutations as effective cancer-specific agnostic biomarkers. Additionally, because ctDNA represents only a small fraction of the total cfDNA, particularly in patients with early-stage tumors (referred to as tumor fraction, TF),^18^ ctDNA detection requires high sensitivity.

Alternatively, SVs are plausible cancer biomarkers. Nevertheless, SVs have not been pursued for liquid biopsy targets as rigorously as mutations and other emerging targets based on epigenetic features, such as DNA methylation and cfDNA fragmentation patterns. SVs often present as DNA copy number alterations (CNA), but the detection of CNA also suffers from very low TF.^19,20^ Tumor-specific translocations can be detected by discordant mate pairs in whole genome sequencing (WGS) data,^21^ which would require deep coverage when applied to cfDNA with low TF. *In silico* selection of short DNA fragments (<150 bp) in WGS libraries could enhance ctDNA signals in cfDNA.^22^ However, our current understanding of ctDNA fragment sizes might be incomplete, as other studies reported that cfDNA fragment sizes in the plasma of cancer patients display more variability, with fragments being both longer and shorter, compared to those in healthy individuals^23,24^. Additionally, tumor-derived high molecular weight DNA in cancer patients’ plasma has also been documented.^25^

The challenges posed by low TF in cfDNA can be addressed by enriching for cancer DNA-specific features through *in vitro* cfDNA processing. To explore this concept, we employed our SV-targeting technology called Genome-wide Analysis of Palindrome Formation (GAPF).^26–30^ The targeted SVs in this case are DNA palindromes, also referred to as inverted repeats or fold-back inversions,^31^ defined by DNA sequences identical to their reverse complements (**Fig. 1A**). While large (>10 kb) germline palindromes are characteristic of sex chromosomes,^32,33^ DNA palindromes in cancer emerge *de novo* through the inverted duplication of single-copy DNA sequences.^26^ Multiple palindromes can form during Breakage-Fusion-Bridge (BFB) cycles that start when a chromosome with two centromeres, known as a dicentric chromosome, forms. Because dicentric chromosomes can result from common adverse events triggering SV generation, such as improper repair of DNA double-strand breaks (DSBs), fusions of dysfunctional and critically short telomeres, and translocations that create chromosomes with two centromeres,^34–36^ DNA palindromes could be widespread in cancer genomes. Notably, anaphase bridges, an indication of segregating dicentric chromosomes, have been observed in preneoplastic regions.^37^ Therefore, DNA palindromes can arise very early during carcinogenesis and may serve as valuable cancer biomarkers in both early and late-stage tumors.

**Fig. 1:**
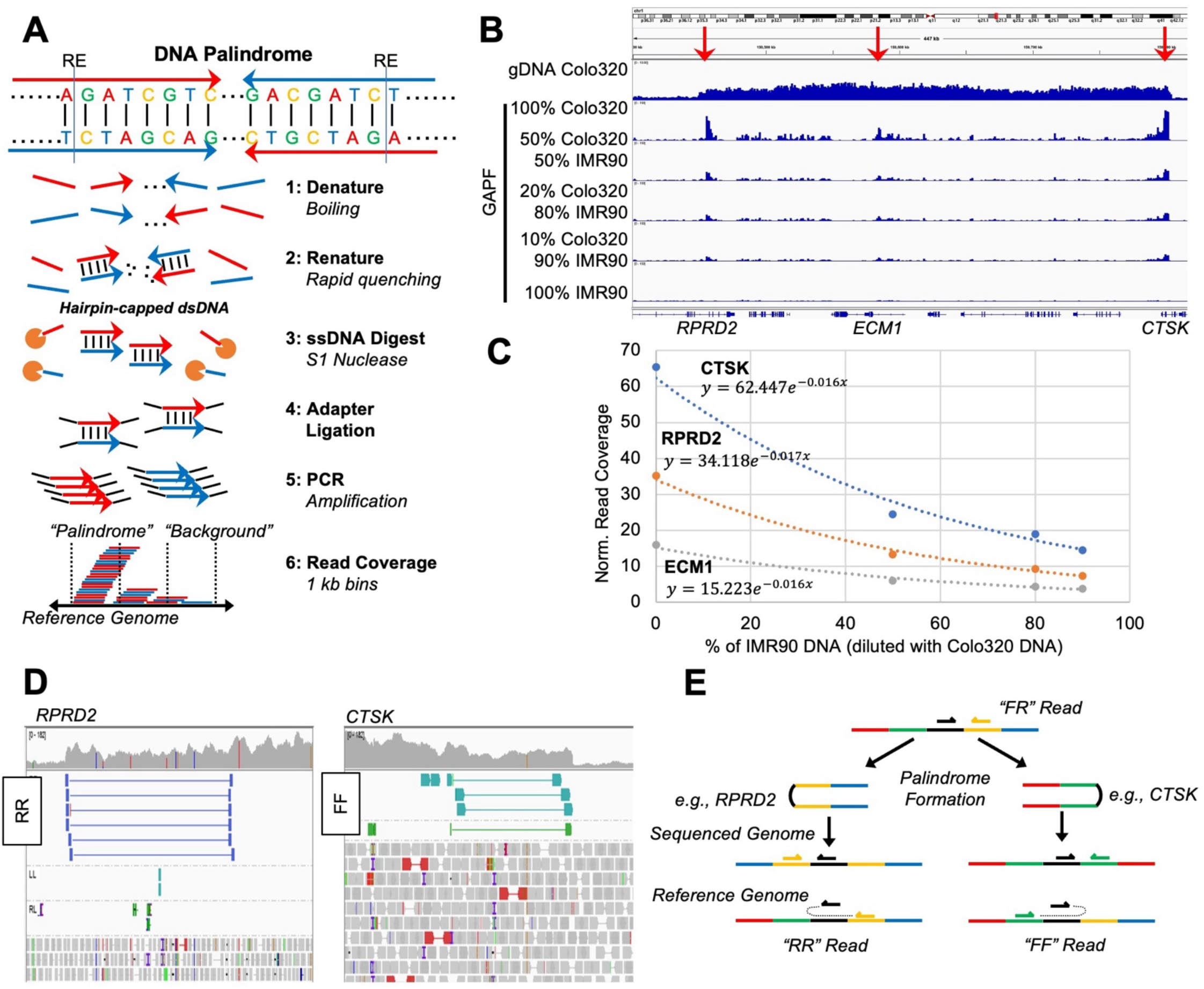
Genome-wide Analysis of Palindrome Formation (GAPF) enriches structural variants. **A)** DNA palindromes are genomic regions with sequences that are reverse compliments of each other and can span large genomic segments. For GAPF, genomic DNA was first digested with restriction enzymes (RE), by which only palindromic junctions folded-back to hairpin-capped dsDNA after denaturing DNA by boiling and rapidly renaturing in an ice water quench. Non-palindromic DNA, including DNA liberated from palindromic junctions by RE, was digested by S1 nuclease. Note that S1 also digested the hairpin part of DNA from palindromic junctions. Following adapter ligation to dsDNA and amplification by PCR was done as a part of NGS library construction. Read coverage was quantified in 1 kb bins genome-wide. **B)** Read coverage in 1 kb bins for genomic or GAPF DNA is visualized using the Integrative Genomics Viewer (IGV). Colo320 colorectal adenocarcinoma cancer cell line DNA is diluted by IMR90 normal fibroblast DNA prior to GAPF. Read coverage in GAPF data show peaks at known palindromic junctions on chr1 at RPRD2, ECM1, and CTSK. **C)** The GAPF read coverage decreases following an exponential trend at RPRD2 (R^2^ = 0.9908), ECM1 (R^2^ = 0.9848), and CTSK (R^2^ = 0.9776) as the amount of diluting IMR90 normal DNA increases. **D)** Paired sequencing reads can have anomalous orientations such as “reverse-reverse” (RR) or “forward-forward” (FF) at sites of structural rearrangements near known palindromic junctions in Colo320 cancer cell line DNA. **E)** A schematic for how palindromes create RR or FF read pairs is presented.

We have previously shown that DNA palindromes can be enriched *in vitro* from genomic DNA using simple molecular procedures of GAPF. Importantly, we have demonstrated palindrome detection in cancer DNA in a 25-fold excess of normal DNA^26^, which underscores the potential of GAPF for cancer detection in samples with low tumor DNA fraction, such as plasma cfDNA. In this study, we applied the principles of GAPF to plasma cfDNA, developing new procedures optimized for small amounts of cfDNA. We also developed a bioinformatics pipeline to help analyze somewhat unique GAPF-seq data structures. By employing traditional machine learning (ML) algorithms for binary classification, we assessed the capability of GAPF-seq to detect cancer DNA in plasma cfDNA, presenting a new tool for liquid biopsy and biomarker discovery. Our findings indicate that GAPF-seq can differentiate cancer patients’ plasma cfDNA from control cfDNA, even in cases of low TF. These results highlight the potential of cancer-specific SVs as sensitive biomarkers in liquid biopsy.

## Results

### Palindrome Detection by GAPF-seq

A new protocol for the Genome-wide Analysis of Palindrome Formation (GAPF) was adapted to address the low levels of ctDNA in plasma cfDNA (**Fig. 1A**). In brief, input genomic DNA (as low as 10 ng) was initially digested with relatively infrequent-cutting enzymes. Subsequently, the DNA was boiled for denaturation and promptly quenched in ice water to facilitate rapid self-annealing of palindromic sequences (i.e., snap-back).^26,38^ The resulting DNA was then digested with the single-strand-specific nuclease S1. The junctions of palindromic DNA could self-anneal following denaturing by boiling, re-forming double-stranded DNA (dsDNA), and becoming resistant to S1. In contrast, non-palindromic sequences would be digested by S1 after denaturation. The remaining dsDNA was ligated with sequencing adaptors and amplified during library construction. The resulting sequencing libraries would be enriched with palindromic junctions.

Approximately 100 million sequencing reads per sample from GAPF-processed DNA were aligned to the hg38 human reference genome and quantified in 1 kb non-overlapping bins (**Fig. 1A**). An accumulation of reads in these 1 kb bins would lead to high coverage, and consequently, high coverage bins (HCBs) may indicate the presence of DNA palindromes.^28,39^ Indeed, when genomic DNA of the Colo320DM cell line was processed by GAPF and sequenced, HCBs were seen in known palindromic junctions on chr1 at *RPRD2*, *ECM1*, and *CTSK* (**Fig. 1B**).^28^ To simulate a low tumor fraction (TF), as expected from plasma cfDNA, Colo320DM DNA was *in vitro* diluted with IMR90 normal fibroblast DNA prior to GAPF. HCBs were consistently detected at *RPRD2*, *ECM1*, and *CTSK* for all dilutions. No such peaks were apparent in the corresponding region of GAPF-seq data from IMR90 DNA (**Fig. 1B**, 100% IMR90). The average normalized read coverage (number of sequencing reads per bin divided by a per-million scaling factor) for HCBs at *CTSK* was 65.5 for 100% Colo320 input DNA and decreased exponentially by dilution (**Fig. 1C**). This exponential trend was similarly observed for *RPRD2* and *ECM1*. The lowest peak was detected at *ECM1* in 10% Colo DNA with an average read coverage of 3.67. Given an average read coverage genome-wide was approximately 0.31, GAPF-seq, even with ultra-low genome-wide coverage, remains capable of reliably detecting palindromic signals after 10-fold dilution.

Paired-end sequencing offers the advantage of utilizing the relative orientation of mate pairs to confirm the presence of palindromes within amplified genomic segments. In WGS data, mates facing the same direction (e.g., forward-forward, FF, and reverse-reverse, RR) relative to the reference genome can be identified at the ends of the amplified region within 1q21. FF or RR read pairs were identified in publicly available deep coverage WGS data (**Fig. 1D**)^40^, but not our shallow WGS, possibly due to the underrepresentation of secondary structure-forming DNA in sequencing libraries. Navigating the secondary structure of palindromic DNA during PCR, which is an integral part of library construction processes, is widely known to be a challenging task.^41,42^

### GAPF-seq for Breast Tumor DNA

To evaluate the ability of GAPF-seq to differentially call tumor DNA from normal DNA, we collected tumor and control GAPF-seq data from 39 pairs of fresh frozen breast tumors and matched leukocytes (buffy coat) (**Fig. 2A**). These breast tumors included both hormone-receptor positive (HR+) and triple-negative breast cancer (TNBC) subtypes, with stages ranging from Stage I to Stage IV. Following GAPF processing, sequencing reads were counted in 1 kb bins genome-wide, and the normalized read coverage was calculated by dividing by a per million scaling factor (**Fig. 1A**). To evaluate the successful enrichment of DNA palindromes, we investigated several quality control measures. First, we examined the read coverage at known inverted repeats on chromosomes 1, 9, 17, 19, and X following GAPF-seq (**Supplementary Table 1**).

**Fig. 2:**
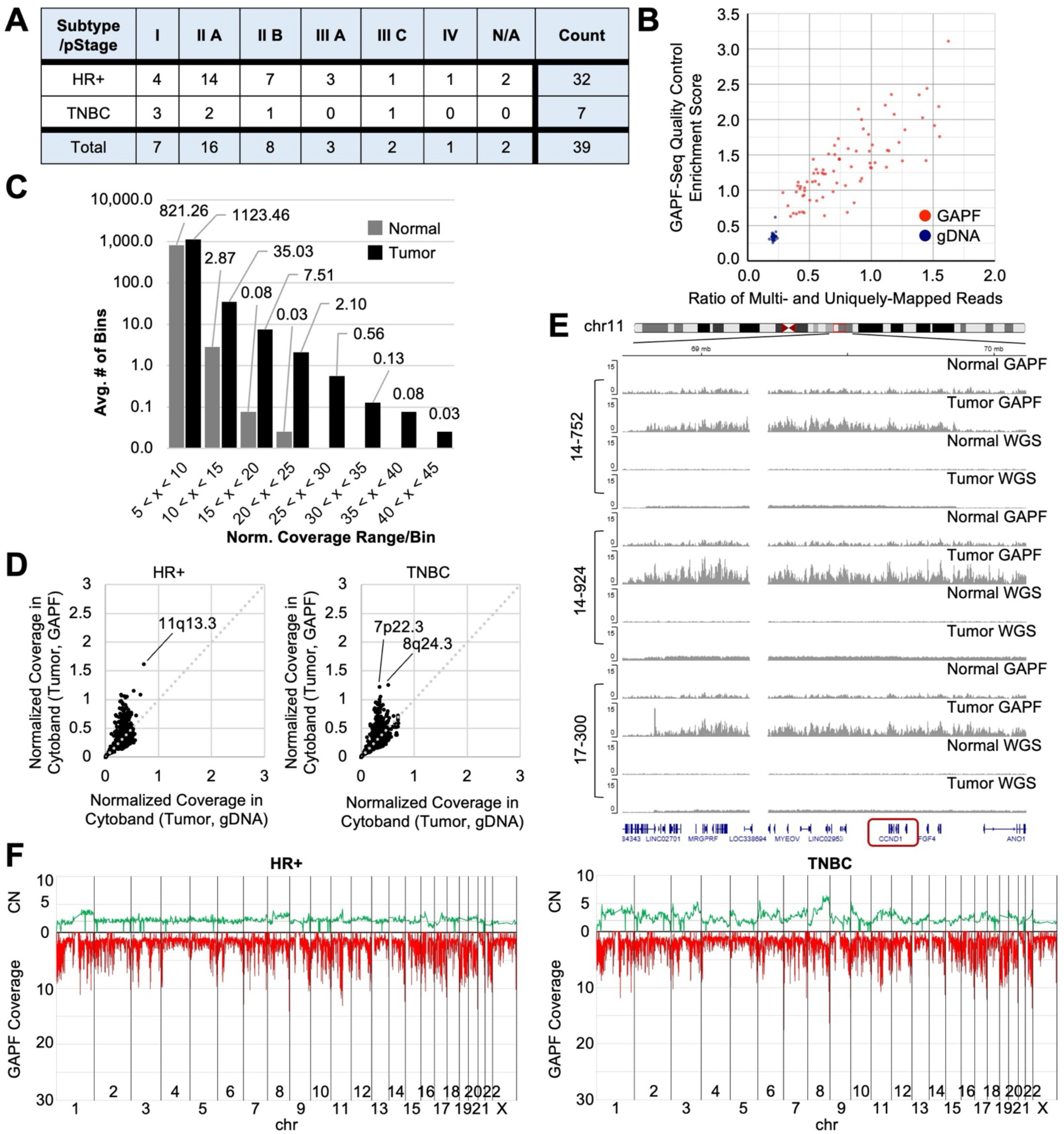
High coverage 1 kb GAPF bins are tumor-specific. **A)** A table showing the breast cancer subtype and pathologic stage (pStage) for 39 breast tumors used for GAPF. **B)** The ratio of multi-mapping to uniquely mapping reads to the reference genome is plotted against the GAPF enrichment score at known palindromic sites. These values are used to evaluate the successful enrichment of DNA palindromes by GAPF. **C)** The average number of 1 kb bins with normalized read coverages in different ranges from 39 breast tumors and matched normal. Out of the 3,209,513 bins in the human genome, the vast majority have very low read coverage in the range between 0 and 5 (not shown). At higher coverage ranges, the average number of bins with high coverage is greater in tumor than normal. **D)** The GAPF read coverage in cytobands is normalized by the size of the cytoband (in bp). The normalized read coverage in each cytoband for genomic DNA (gDNA) is plotted against GAPF DNA and averaged for hormone receptor-positive (HR+) breast tumors (left, n = 26) or triple negative breast cancer (TNBC) (right, n = 5) tumors. **E)** Read coverage in 1 kb bins for genomic (WGS) or GAPF DNA is visualized using the Integrative Genomics Viewer (IGV) in select HR+ breast tumors at a region on chr11 near *CCND1*. The read coverage with GAPF is significantly more pronounced than by WGS. **F)** The average copy number (CN) is plotted against the average GAPF read coverage for HR+ breast tumors (top, n = 26) and TNBC tumors (bottom, n = 5).

Enrichment of these germline inverted repeats would indicate that GAPF-seq would be similarly successful at enriching *de novo* DNA palindromes elsewhere in the genome. The normalized read counts for each of the control regions were then averaged to derive an enrichment score. Because the average normalized read coverage genome-wide is approximately 0.3, GAPF-seq was considered successful at amplifying DNA palindromes with an enrichment score above that while samples with scores below 0.4 were excluded from the analysis. Another metric for evaluating GAPF-seq is the ratio of concordant read pairs mapped more than once (multi-mapped) to read pairs mapped only once (uniquely-mapped). During GAPF processing, DNA is denatured and renatured so strands with complimentary sequences located nearby, such as DNA palindromes, but also other types of DNA repeats that are abundant in the genome, are enriched. This results in a majority of sequencing reads originating from repetitive regions of the genome and having multiple possible alignments. The ratio of multi- and uniquely-mapped read pairs in genomic DNA not processed by GAPF is approximately 0.25 whereas GAPF-seq DNA can have ratios much greater than 1.0. Plotting the GAPF-seq quality control enrichment score against the ratio of multi- and uniquely-mapped reads visualizes the distinct alignment and mapping characteristics of DNA processed by GAPF (**Fig. 2B**). To test the reproducibility of GAPF-seq, we assessed the average read coverage of uniquely-mapped reads in each cytoband (instead of 1 kb bins) throughout the genome using GAPF-seq replicates from seven pairs of tumor and buffy coat DNA (**Supplementary Fig. 1**). The read coverage at the cytoband level showed remarkable consistency, with an intraclass correlation coefficient of 0.964 (95% CI: 0.962-0.966), confirming the assay’s reproducibility.

A vast majority of 1 kb bins (approximately 3,130,000 bins out of 3,209,513 total bins) in both breast tumor and buffy coat DNA exhibited an average uniquely mapped read coverage of less than 5, likely attributed to the digestion of non-palindromic sequences by S1 nuclease. However, there was a notable difference in the number of 1 kb bins with read coverage >5 between tumor and control samples (**Fig. 2C**). Particularly at higher coverage ranges (read coverage >10), tumor GAPF-seq data displayed a greater number of bins than control GAPF-seq data, indicating the potential presence of tumor-specific palindromes. This observation formed the basis for delineating HCBs as means to differentiate tumor and normal DNA.

At the cytoband level, notable enrichment of GAPF read coverage over WGS read coverage was seen in a subtype-specific manner, with 11q13.3 enrichment in HR+ tumor DNA and 7p22.3 and 8q24.3 in TNBC tumor DNA (**Fig. 2D**). The cytoband 11q13.3 houses the breast cancer oncogene *CCND1*, which encodes the cyclin D1 protein involved in cell cycle progression and is frequently amplified in HR+ breast cancer subtypes. The read coverage around *CCND1* shows dramatic enrichment with GAPF-seq compared to shallow WGS in breast tumors with amplification in this region (**Fig. 2E**).

As DNA palindromes are frequently associated with genomic amplification, the average copy number (CN) of chromosomal segments from shallow WGS data was plotted against the read coverage of GAPF-seq data for 26 HR+ and 5 TNBC tumors (**Fig. 2F**). Overall, TNBC showed more CNA than HR+ breast tumors, aligning with previous reports.^43^ While genome-wide GAPF read coverage appeared indistinguishable between subtypes, a closer look at GAPF-seq and WGS data from individual pairs revealed unique subtype coverage patterns. For example, patient 14-990 with TNBC exhibited significant genome-wide CNA (**Supplementary Fig. 2**), and GAPF-seq revealed a notable read enrichment in cytoband 8q24.3 that correlated with increased CN. In contrast, a patient with HR+ breast cancer (14-752) showed CNA and increased GAPF coverage localized primarily to cytoband 11q13 (**Supplementary Fig. 2**). Accordingly, cytobands 11q13.2-5 were enriched in HR+ tumors, while cytoband 8q24.3 was predominant in TNBC.

### Tumor DNA Calling Threshold by GAPF-seq

Upon further inspection of regions of high GAPF coverage, we observed common enrichment in several bins across both tumor and control samples (**Supplementary Fig. 3**). We explored the possibility of germline palindromes.^44,45^ Alternatively, we considered that specific DNA segments were resistant to denaturation, such as segments of high GC content, and consequently, these segments would be resistant to S1 nuclease, leading to amplification during library construction. This phenomenon could result in non-palindromic enrichment during GAPF.^38^ Because of these technical issues, we excluded 1 kb bins consistently enriched in both tumor and normal GAPF-seq samples (75,765 bins, ~2.5% of the entire set of 3,209,513 bins) to focus on identifying tumor-specific *de novo* DNA palindromes (**Supplementary Table 2**).

The remaining 3,133,748 bins were then arranged in descending order based on their read depth coverage (**Fig. 3A**). The observation that HCBs were tumor-specific (**Fig. 2C**) prompted us to investigate differential calling between tumor and normal DNA using the chromosomal distribution of the top 1,000 HCBs, visualized as GAPF profiles in radial plots (**Fig. 3B**). The examination of normal DNA GAPF profiles revealed HCBs across all chromosomes (**Fig. 3B, Supplementary Table 3**). In contrast, tumor GAPF profiles exhibited HCBs primarily originating from one or a few chromosomes, suggesting the potential clustering of *de novo* palindromes in the breast tumor DNA. The nearly uniform distribution of HCBs across all chromosomes in the normal GAPF profiles suggests the randomness of residual dsDNA after denaturation that were amplified through GAPF, as expected in a sample lacking *de novo* palindromes in single-copy (unique) regions of the genome. We also confirmed that GAPF profiles were highly reproducible (**Fig. 3C, Supplementary Table 4**). The noticeable tumor-specific clustering of HCBs on chromosomes hinted at the intriguing possibility of identifying a global threshold for classifying samples as tumor or normal (threshold model).

**Fig. 3.**
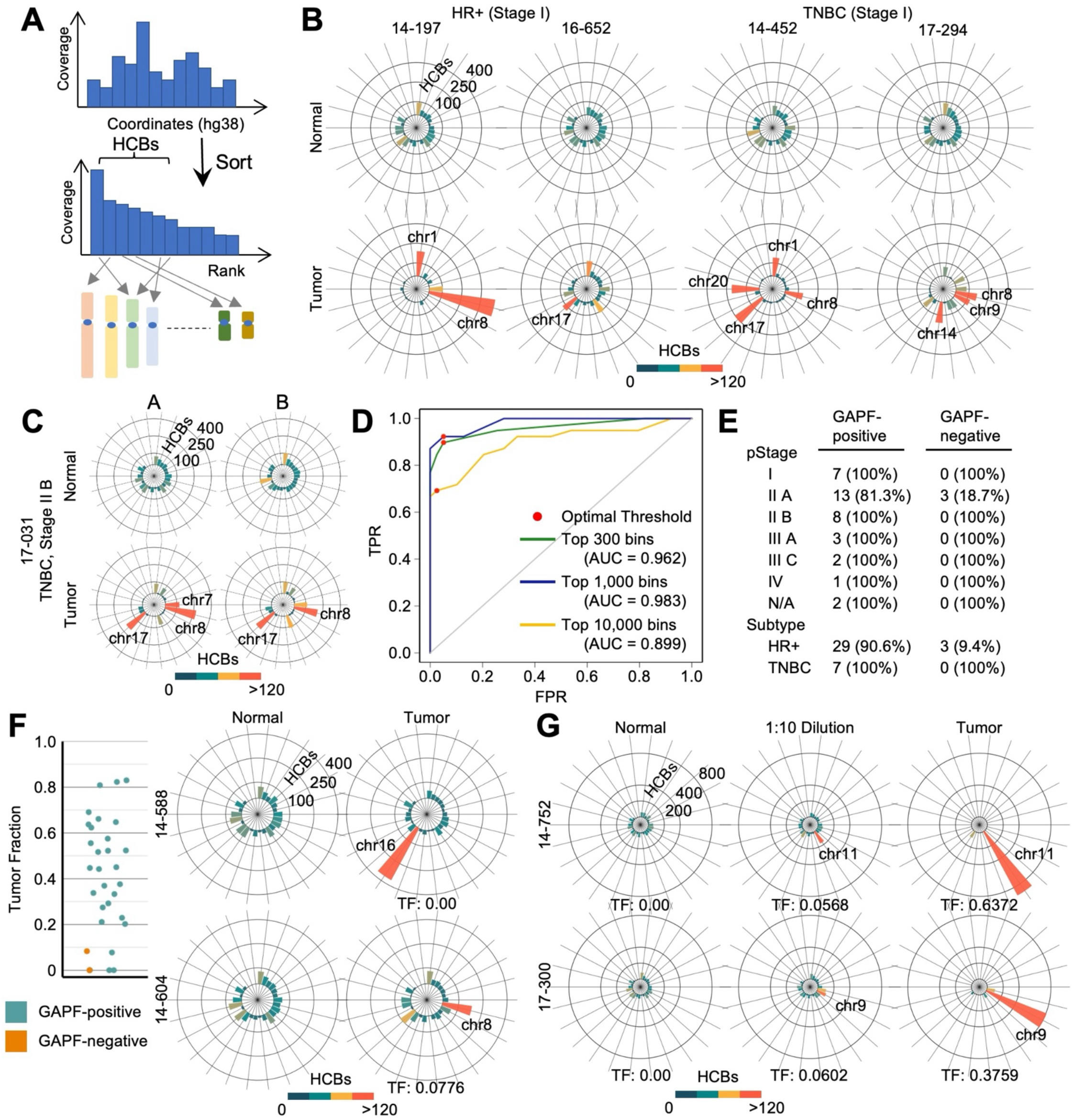
High coverage GAPF bins classifies tumor or normal DNA using a threshold model. **A)** The analytical approach for GAPF-seq sorts 1 kb bins in descending order according to the normalized read coverage and identifies the chromosome of origin for the high coverage bins (HCBs) included in subsequent analyses. **B)** GAPF profiles for breast tumors and matched normal buffy coat visualized in radial charts show clustering of the top 1,000 HCBs on chromosomes. **C)** GAPF profiles using the top 1,000 HCBs are reproducible as visualized in radial charts for two breast tumor pairs performed in duplicate. **D)** Receiver operating characteristic (ROC) curves generated by varying the threshold for the number of HCBs on a chromosome as the classification criterion for tumor or normal DNA. The top 300 (green), 1,000 (blue), or 10,000 (yellow) HCBs were considered. The optimal threshold (red dot) is determined by Youden’s J statistic which maximizes sensitivity and specificity. **E)** GAPF classification results for 39 breast tumors by stage using a chromosome threshold of 100 HCBs. **F)** Dot plot showing the tumor fraction (TF) by GAPF classification (blue = GAPF-positive, orange = GAPF-negative). n = 19. The GAPF radial plot for two breast tumors with TF = 0 show enrichment by GAPF-seq. **G)** GAPF profiles for normal DNA (left column), tumor DNA diluted 10x by normal DNA (middle), and tumor DNA (right).

To assess the optimal number of HCBs for effectively differentiate tumor and normal DNA, we generated receiver operating characteristic (ROC) curves using the top 300, 1,000, or 10,000 HCBs and evaluated their chromosomal distribution (**Fig. 3D, Supplementary Table 5 and 6**). The classification criterion for determining whether a sample contains tumor DNA (GAPF-positive) or not (GAPF-negative) relied on whether HCBs exceeded a global threshold on any single chromosome. For the top 1,000 bins, the global threshold tested ranged from 60, classifying every sample as GAPF-positive, to 915, classifying every sample as GAPF-negative. With three-fold cross-validation and three different random assignment seeds, the average area under the ROC curve (AUC) for the binary classifier using the top 1,000 HCBs was 0.9829 (s.d. ± 0.0080). For each cross-validation fold for each seed, an optimal global threshold was then determined using Youden’s J statistic, aiming to maximize the sensitivity and specificity of the binary classifier and achieving an average sensitivity of 87.18% (s.d. ± 9.42%) and specificity of 95.73% (s.d. ± 4.05%). Using the top 300 or 10,000 HCBs instead yielded AUC values of 0.9624 (s.d. ± 0.0118) and 0.8992 (s.d. ± 0.0332), respectively, indicating the top 1,000 as the most effective HCB group for the binary classifier in this dataset.

Using the top 1,000 HCBs and the best performing threshold model, we identified 36 GAPF profiles with at least one chromosome above the 105-bin threshold (GAPF-positive), while 3 tumors were misclassified as normal (GAPF-negative) (**Fig. 3E**). Notably, the clustering of HCBs on a chromosome was evident for all stage I tumors (TNBC or HR+), demonstrating that GAPF-seq could detect early-stage breast cancers (**Fig. 3B**, **Fig. 3E**). The three GAPF-negative samples were all HR+ breast tumors. HR+ tumors typically exhibit fewer SVs than TNBC,^43^ which may explain these GAPF-negative cases. As SVs often manifest as CNA, we quantified CNA for 31 breast tumors using shallow WGS and ichorCNA, a tool for estimating tumor fraction (TF) in plasma cell-free DNA (**Fig. 3F**). GAPF-negative tumors showed low CNA; however, several samples with low CNA were GAPF-positive (14-604 and 14-588). To further examine whether GAPF-seq could call tumor DNA for tumors with very low CNA, we ran GAPF-seq with 10-fold *in vitro* dilutions of tumor DNA with paired normal DNA. All samples yielded the expected tumor DNA content after 10-fold dilution of tumor DNA. Based on the threshold model (**Fig. 3D**), seven out of 8 samples were accurately classified as tumor DNA, indicating that GAPF profiles have the potential to identify tumors with low DNA content (**Fig. 3G, Supplementary Table 7**).

### Subtype-specific Clustering of High Coverage Bins

By examining chromosomes frequently carrying top 1,000 HCBs above the threshold, chr1 and chr8 appeared to be common between HR+ and TNBC subtypes, while chr16 and chr11 were unique to HR+ and chr7 was unique to TNBC (**Fig. 4A**). We further investigated HCBs at the cytoband level, comparing the number of GAPF top 1,000 HCBs in each cytoband from tumor GAPF profiles to that of normal GAPF profiles (**Fig. 4B**). In TNBC, on average 197.6 of 1,000 HCBs were in 8q24.3, the telomeric end of 8q, in contrast to 28.0 HCBs in normal DNA. The 8q24.3 cytoband was also a hotspot for enrichment in HR+ tumor DNA and, to the lesser extent, in normal DNA (**Fig. 4C**). HCBs were often clustered in cytobands close to telomeres. In addition to 8q24.3, clustering was observed in the telomeric cytobands 13q34 (in TNBC) and 16p13.3 (in HR+) in a tumor subtype-specific manner (**Fig. 4B**, **Fig. 4C**). The fusions of critically short telomeres followed by BFB cycles^46^ may underlie the enrichment of palindromes and HCBs at the cytobands of chromosome ends. Notable exceptions include cytobands 8q24.21 (in TNBC) and 11q13.2-5 (in HR+), which house prevalent breast cancer oncogenes *MYC* and *CCND1*, respectively, and are frequently amplified. Indeed, the amplification of the *CCND1* locus was strongly associated with HCBs (**Fig. 4D**). Complex CNA patterns at 11q13 were common in HR+ tumors, with another amplicon telomeric to the *CCND1* amplicon associated with HCBs.

**Fig. 4:**
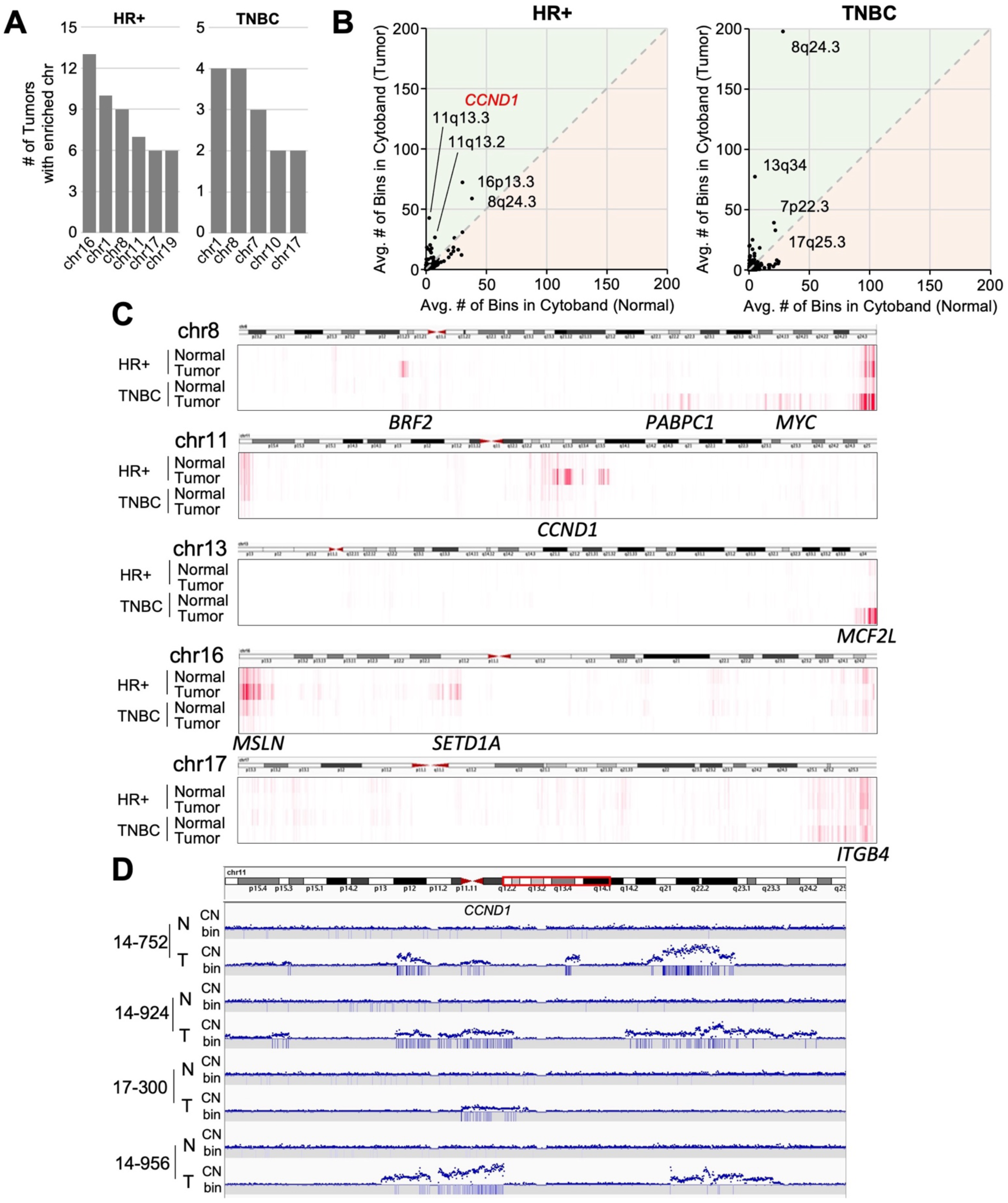
Clustering of high coverage bins is breast cancer subtype-specific. **A)** The number of tumors (out of 39) with chromosomes enriched with high coverage bins (HCBs) above the threshold in Fig. 3D for HR+ and TNBC tumors. **B)** The average number of HCBs in each cytoband in tumors is plotted against the number of HCBs in each cytoband in normal samples. Points above the diagonal indicate enrichment of the cytoband by GAPF. **C)** A heatmap showing the location of HCBs on chr8, chr11, chr13, chr16, and chr17 for HR+ and TNBC tumor and normal DNA. **D)** CN and HCBs (bin) for four (4) HR+ breast tumors near CCND1.

Aberrant read pairs spanning palindromic junctions are likely underrepresented,^41,42^ and deep coverage would be necessary to identify FF or RR read pairs (**Fig. 1D**). Nonetheless, we identified multiple RR read pairs at the copy number breakpoint of the amplicon at 11q13.2 with HCBs clustering nearby (**Supplementary Fig. 4**).

### Machine Learning Algorithms for Binary Classification

Aside from the Youden’s J statistic-based threshold model for our dichotomous diagnostic test, a variety of statistical, probabilistic, and optimization techniques can be applied to genomic data using machine learning (ML) for binary classification. Traditional ML models can be quickly adapted when test and control datasets have the same number of features, such as in GAPF profiles, where both tumor and normal GAPF profiles have 23 features.^47^ An automated end-to-end ML pipeline called STREAMLINE^48^ was used for the binary classification of the breast tumor and paired normal DNA (as in **Fig. 3**). The number of top 1,000 HCBs on each chromosome from the GAPF-seq data was input as a two-dimensional array, and STREAMLINE attempted to identify the samples as tumor or normal DNA. STREAMLINE, a recently developed autoML tool, facilitates ML modeling by enforcing rigorous modeling algorithm comparisons, analysis transparency, reproducibility, and sensitivity to complex associations in data. In the first year of its release, it has been successfully applied to various biomedical ML modeling and data mining tasks.^49,50^ After basic data cleaning and three-fold cross-validation partitioning (**Fig. 5A**), ML models were trained using Naïve-Bayes (NB), random forest (RF), extreme gradient boosting (XGB), category gradient boosting (CGB), and extended supervised tracking and classifying system (ExSTraCS) algorithms. With a random sample assignment seed of 42, all models yielded very high average AUC values (**Fig. 5B**). To further mitigate batch effects from partitioning, ML modeling was repeated using two additional random assignment seeds (22 and 32), and the AUC results from all cross-validation folds from the three seeds were averaged (**Fig. 5C**). Even with this rigorous validation by ML modeling, average AUC values were greater than 0.925 for all models, with the Naïve-Bayes model demonstrating the highest performance (Avg. AUC = 0.957). Each unique seed yielded a predicted class for a sample, and a heatmap visualized the congruence of true positive outcomes from the testing sets of the models generated by three separate random assignment seeds. Binary classification was generally consistent among models, with a few exceptions (**Fig. 5D**).

**Fig. 5:**
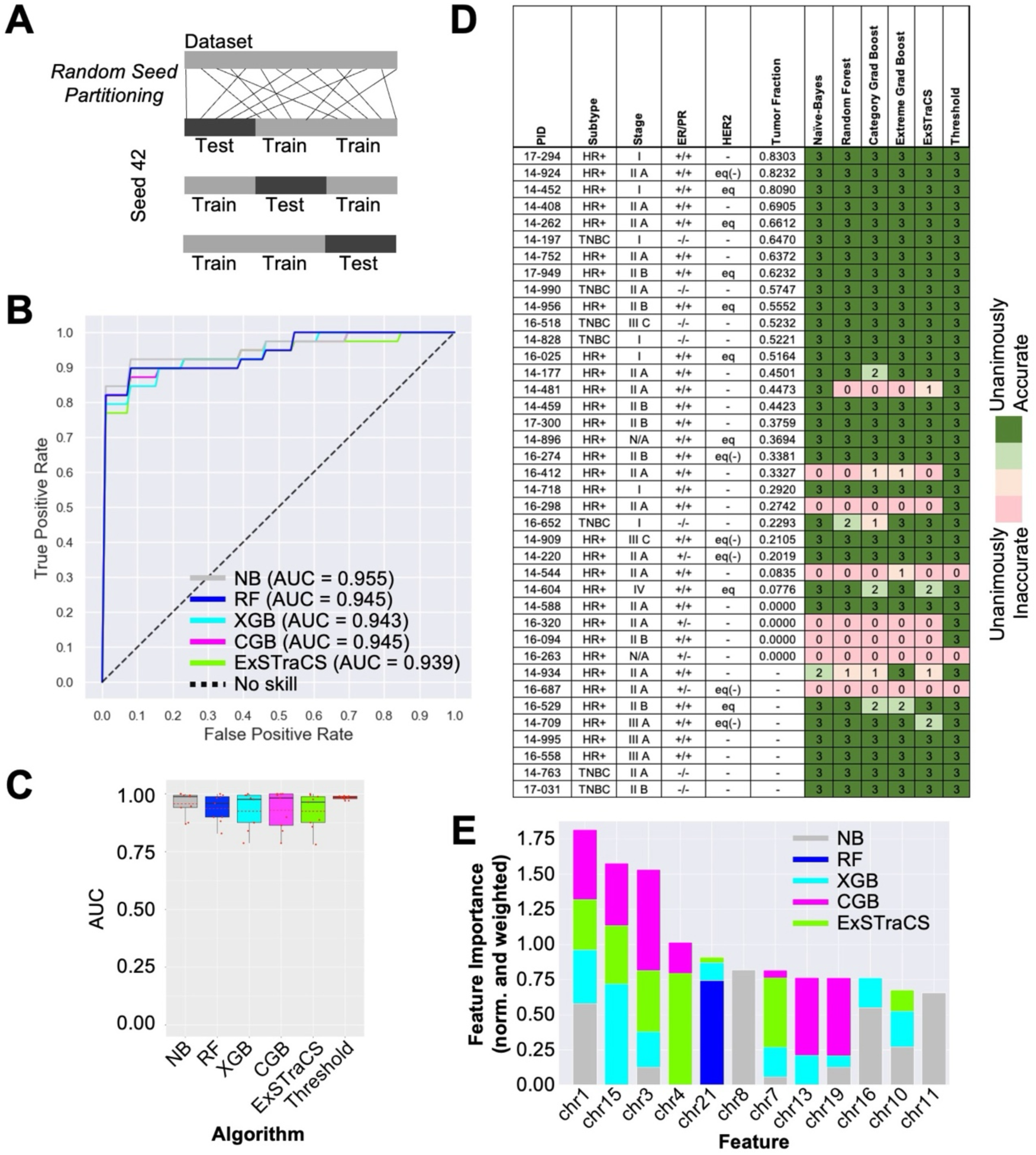
A traditional machine learning pipeline with GAPF-seq accurately classifies tumor and normal DNA. **A)** Random seed partitioning of the dataset assigns data points into groups for cross-validation (CV) testing of modeling. For three-fold CV, two groups are used to train the model while the remaining group is used to test the model. **B)** ROC curves for machine learning (ML) models generated by STREAMLINE classifying tumor or normal DNA using the Naïve-Bayes (NB), random forest (RF), extreme gradient boosting (XGB), category gradient boosting (CGB), and extended supervised tracking and classifying system (ExSTraCS) algorithms with a random assignment seed of 42 and three-fold CV. **C)** Box plots showing the AUC from each of 3 cross-validation testing sets for models generated using random assignment seeds of 22, 32, and 42. Red dot = AUC value, red dotted line = mean AUC. **D)** Table of breast tumor subtype, stage, ER/PR/HER2 status, and tumor fraction (TF) with a heatmap showing the congruence of classification results from random assignment seeds and the threshold and machine learning (ML) models. **E)** Composite normalized and weighted feature importance bar plots. Scores are normalized within each algorithm and weighted by the median AUC for that algorithm. Composite importance bar plots normalize feature importance scores within each algorithm and then weights these normalized scores by the median AUC for that algorithm to consider accurate models more than less accurate ones.

Traditional models can provide clues about which features contribute most to a prediction.^47^ The STREAMLINE ML pipeline also provided the estimation of the relative importance of each chromosome in accurately predicting tumor or normal DNA (**Fig. 5E**). Feature importance was determined by randomly permutating one chromosome at a time in the testing data and evaluating its impact on the performance of each ML model.^51^ Interestingly, non-linear classification models RF, XGB, CGB, and ExSTraCS deemed chromosomes such as chr15, chr3, chr4, and chr21 as important, which contrasts with our threshold model (**Fig. 4A**). These chromosomes exhibited depletion of HCBs in tumor GAPF profiles relative to normal profiles. For example, normal samples had an average of 20.1 of the top 1,000 HCBs on chr15, whereas tumor tissue had only 9.6. In contrast, for chromosomes frequently carrying HCBs above the threshold, such as chr8 and chr11, the average numbers of HCBs were 48.9 and 64.5 in normal samples, and 131.1 and 118.5 in tumor samples, respectively. Only the linear classifier algorithm Naïve-Bayes yielded high feature importance scores for chr1, chr8, and chr11, all of which were frequently enriched with HCBs above the threshold determined by Youden’s J statistic.

### Long cfDNA Fragments and Tumor Fraction

With the ability to differentiate tumor DNA from paired normal DNA, we proceeded to evaluate GAPF-seq using plasma-borne cfDNA. GAPF-seq presumably relies on sufficiently long DNA fragments to ensure that both arms of DNA palindromes are physically tethered together in a continuous fragment, facilitating intra-strand annealing by snap-back. However, prevalent methods for extracting cfDNA from plasma employ affinity-columns or -beads and predominantly yield fragments bound to mono- or di-nucleosomes, which would be too short to capture palindromes. Whether very long circulating tumor DNA (ctDNA) fragments exist in the plasma of cancer patients remains a topic of debate.^52^ To investigate whether long cfDNA fragments exist and are as feasible for cancer detection as short cfDNA fragments, we isolated cfDNA using 0.5 mL plasma from 30 advanced prostate cancer (PCa) patients using either affinity-beads (Bead) or traditional phenol-chloroform extraction (PCE) (**Fig. 6A**). Plasma from advanced prostate cancer (PCa) patient is expected to carry sufficient ctDNA for analyses whereas the majority of our breast tumor samples are Stage II or earlier and less likely to have high amounts of ctDNA in plasma. Additionally, the use of PCa cfDNA serves as an initial test for GAPF-seq as a pan-cancer approach. The bead-based cfDNA isolation kit yielded an average of 7.21 ng of cfDNA per 500 μL of plasma, with only 4 samples yielding more than 10 ng. In contrast, PCE yielded 24.02 ng on average with 17 samples yielding more than 10 ng. Due to the low amount of cfDNA extracted by the bead-based isolation kit, there was often insufficient cfDNA for additional characterization experiments beyond sequencing. Furthermore, while the bead-based cfDNA isolation kit yielded mono-, di-, and even some tri-nucleosomal cfDNA, PCE extraction resulted in a broad range of cfDNA fragment sizes, including nucleosome-sized fragments (**Fig. 6B**) and, notably, fragments longer than 10 kb, which would be amenable to snap-back (**Fig. 6C**). We then used fragmentation and NGS library construction for shallow WGS to evaluate TF using ichorCNA. CNA patterns appeared similar between affinity bead-extracted and PCE cfDNA (**Fig. 6D**). The TF in PCE cfDNA showed remarkable consistency with that in bead-extracted cfDNA with an intraclass correlation coefficient of 0.988 (95% CI: 0.967-0.996) (**Fig. 6E**), demonstrating that PCE cfDNA contains tumor DNA. To examine whether or not the long fragment population of extracted cfDNA is contributing to the overall TF, we evaluated TF for paired sequencing reads with short (<200 bp) and long (≥200 bp) insert sizes (**Supplementary Fig. 5**), assuming that read pairs with >200 bp insert sizes likely originated from long DNA fragments after fragmentation rather than mononucleosomal fragments. TF was equally represented in both populations of size-selected reads, indicating that long cfDNA fragments would also be feasible for cancer detection.

**Fig. 6.**
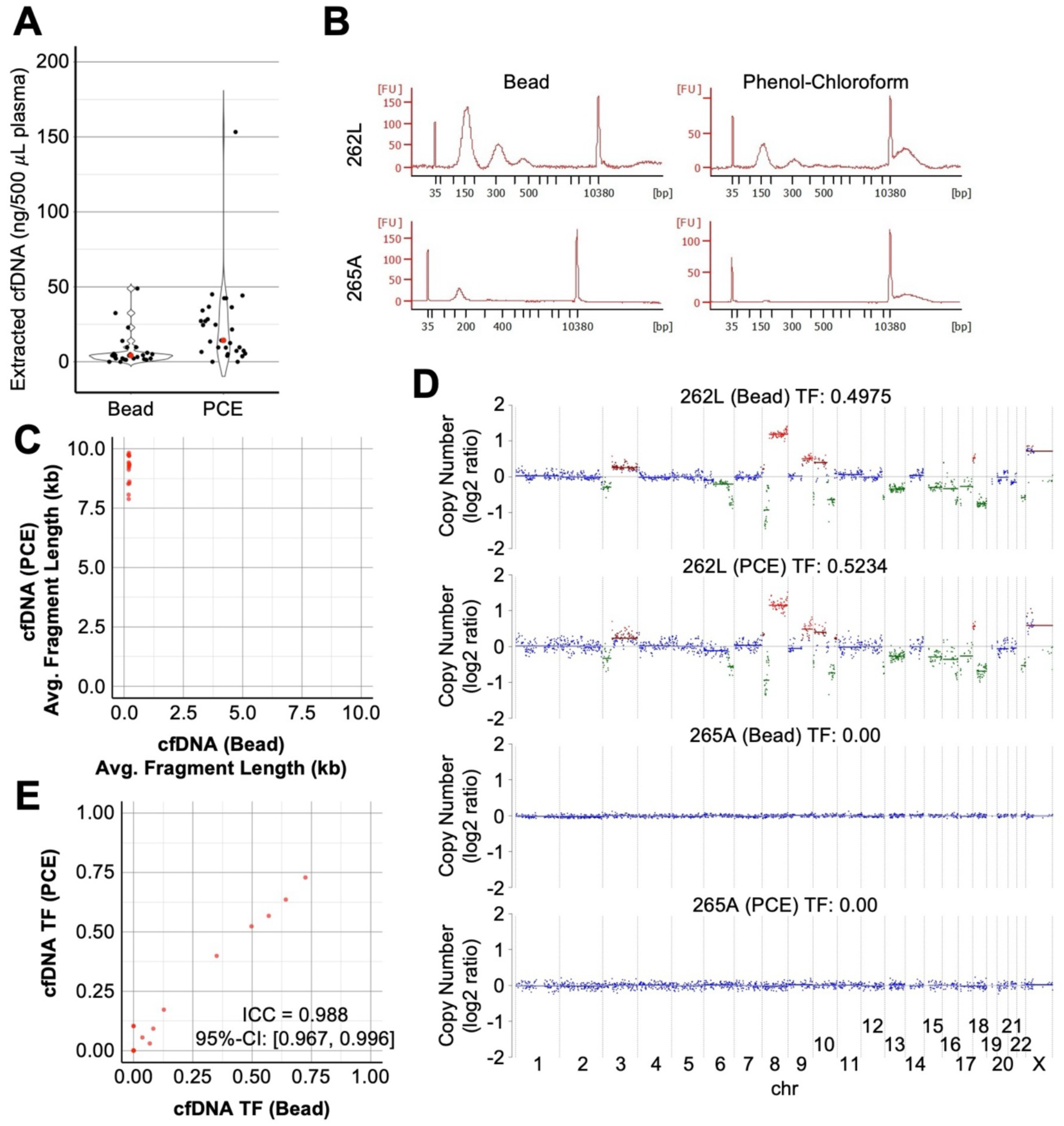
Long fragment cell-free DNA (cfDNA) extracted from plasma contains circulating-tumor DNA (ctDNA). **A)** The amount of cell-free DNA (cfDNA) extracted from prostate cancer (PCa) patients using affinity-bead based extraction (bead) (n = 22) or phenol-chloroform extraction (PCE) (n = 30) normalized to 0.5 mL plasma. Red dot = mean. **B)** Fragment analysis of cfDNA isolated from the blood of PCa patients using affinity-bead based extraction (bead) or phenol-chloroform extraction (PCE). **C)** The average fragment length of plasma cfDNA extracted using affinity-beads is plotted against that using phenol-chloroform. **D)** Copy number (CN) and tumor fraction (TF) of cfDNA extracted using affinity beads (bead) or phenol-chlorofom (PCE) as calculated by ichorCNA. **E)** The TF of cfDNA extracted using affinity-beads (bead) is plotted against that using phenol-chloroform (PCE). The intraclass correlation coefficient is 0.998 with a 95% CI of 0.967 to 0.996.

### High Coverage Bins on chrX as a Determinant of Binary Classification

GAPF profiles with the top 1,000 HCBs were generated from cfDNA from 30 PCa patients. Matched leukocyte DNA was available from 10 of the PCa patients to generate GAPF profiles (**Fig. 7A, Supplementary Table 8**). Similar to the GAPF profiles from breast tumor and normal DNA, HCBs from normal DNA were distributed across every chromosome. In contrast, PCa cfDNA exhibited modest clustering of HCBs. Given that plasma cfDNA and DNA extracted from the buffy coat underwent different pre-analytical steps, it is essential to address whether the binary classification results could potentially be influenced by differences in sample processing. To investigate this issue, GAPF-seq was performed on 24 plasma samples obtained from age-matched males without any known diseases (**Supplementary Table 9**).

**Fig. 7.**
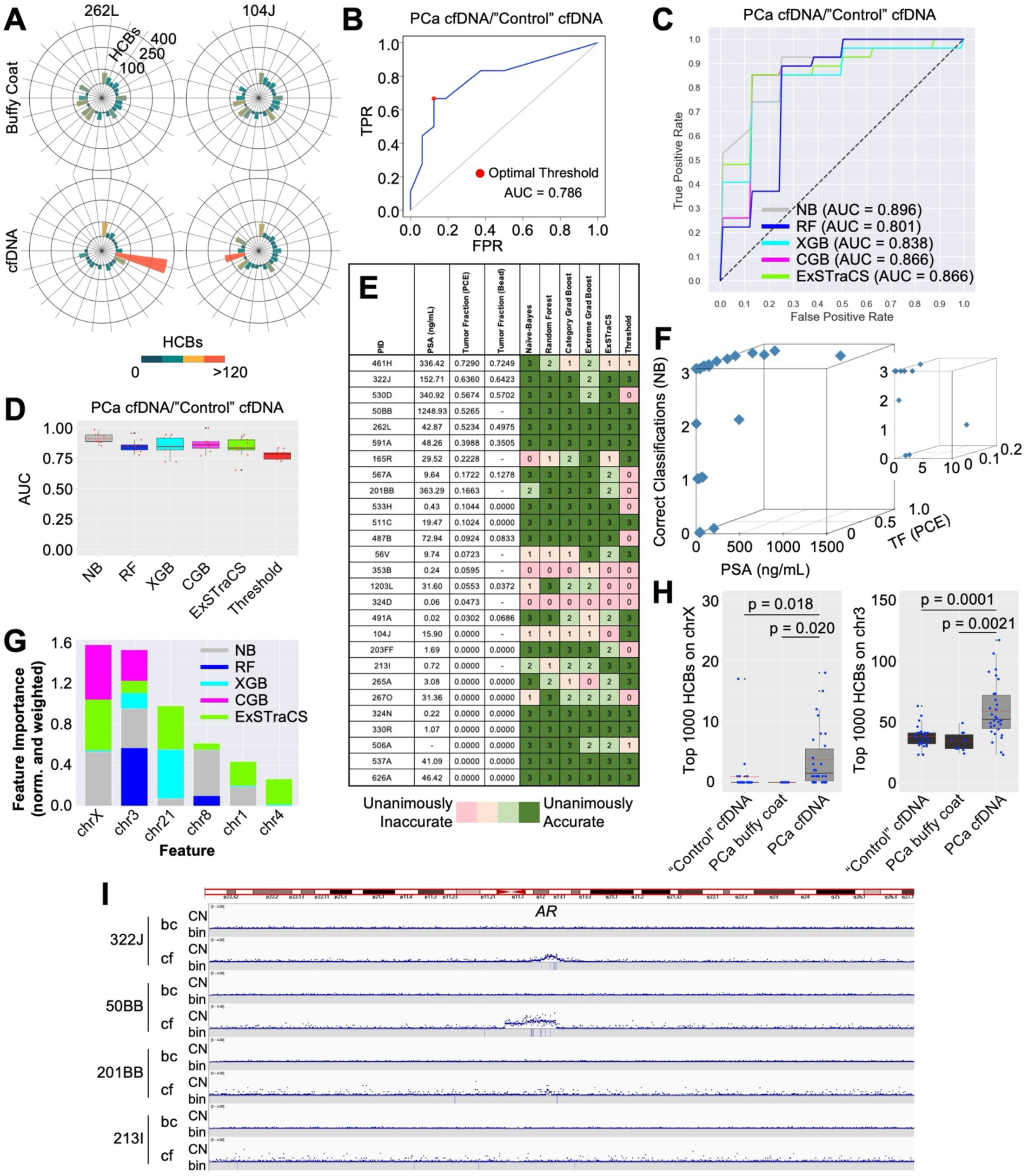
GAPF-seq accurately classifies PCa cfDNA and points to chrX as a key determinant factor. **A)** GAPF profiles using the top 1,000 high coverage bins (HCBs) of prostate cancer (PCa) cfDNA and matched buffy coat DNA visualized in radial plots. **B)** ROC curve generated by varying the threshold for the number of top 1,000 HCBs on a chromosome as the classification criterion for PCa cfDNA or age- and gender-matched cfDNA extracted from individuals with no known disease. The optimal threshold (red dot) is determined by Youden’s J statistic. **C)** ROC curves for machine learning (ML) models generated by STREAMLINE classifying tumor or normal DNA using the Naïve-Bayes (NB), random forest (RF), extreme gradient boosting (XGB), category gradient boosting (CGB), and extended supervised tracking and classifying system (ExSTraCS) algorithms with a random assignment seed of 42 and three-fold CV. **D)** Box plots of AUC values for classifying PCa cfDNA and control cfDNA for machine learning (ML) classifiers using the NB, RF, XGB, CGB, or ExSTraCS algorithms with three-fold CV and random assignment seeds of 22, 32, and 42. Red dot = AUC value, red dotted line = mean AUC. **E)** Table of PSA (ng/mL), tumor fraction (TF) calculated by ichorCNA from cfDNA extracted by PCE or affinity-beads with a heatmap showing the congruence of classification results from random assignment seeds of 22, 32, and 42 and different classification models. **F)** A 3D plot showing PSA (ng/mL), tumor fraction (TF) as calculated from phenol-chloroform extracted cfDNA, and the number of correct classifications by the Naïve-Bayes algorithm across random assignment seeds of 22, 32, and 42. **G)** Composite normalized and weighted feature importance bar plots. Scores are normalized within each algorithm and weighted by the median AUC for that algorithm. **H)** Box plots showing the number of top 1,000 HCBs on chrX or chr3 for GAPF-seq data using age- and gender-matched cfDNA (n = 11), buffy coat DNA from PCa patients (n = 10), and cfDNA from PCa patients (n = 27). Red dot = AUC value, red dotted line = mean AUC. **I)** CN and HCBs (bin) for PCa cfDNA on chrX near AR

These control plasma samples were classified alongside the 27 plasma samples from PCa patients that passed GAPF-seq quality control using ML modeling. An ROC curve using the top 1,000 HCBs from 27 PCa patients and 24 controls showed that the average AUC was 0.781 (s.d. ± 0.039) (**Fig. 7B**). The optimal global threshold determined by Youden’s J statistic was 105 bins, resulting in a sensitivity of 58.03% (s.d. ± 14.46%) and a specificity of 83.33% (s.d. ± 13.98%). STREAMLINE was then used to generate binary classification models (**Fig. 7C**). The AUC values from each cross-validation set from 3 different random assignment seeds were averaged together (**Fig. 7D**). Consistent with the breast tumor/normal DNA classification (**Fig. 5C**), the best-performing model, on average, utilized the NB algorithm (Avg. AUC = 0.896).

TF values were grouped based on this output from GAPF profiles by our threshold model (**Fig. 7C**). A heatmap visualized the congruence of true positive outcomes, from unanimously accurate to unanimously inaccurate, against the prostate-specific antigen (PSA) level at the time of blood draw, and TF of PCa cfDNA extracted by phenol-chloroform or affinity-beads (**Fig. 7E**). The threshold model failed to consistently classify PCa cfDNA correctly with 5 samples with TF>0.1 (461H, 530D, 567A, 201BB, and 533H), the cutoff that ichorCNA indicates the presence of ctDNA with high confidence, being misclassified as GAPF-negative, although most of these cases were classified correctly by ML modeling. On the other hand, 9 samples with an estimated TF of zero (104J, 203FF, 213I, 265A, 267O, 324N, 330R, 506A, 537A, and 626A) were correctly classified as GAPF-positive by most of the models. Importantly, four cases exhibited normal PSA levels, suggesting that GAPF-seq might detect ctDNA even when clinical signs are mild. A three-dimensional plot of the number of correct classifications from the three different Naïve-Bayes models generated by the random assignment seeds against the PSA level and TF of cfDNA (PCE) shows that PSA and TF are not necessarily correlated with increases in classification accuracy (**Fig. 7F**). Overall, Naïve-Bayes models were unanimously accurate in identifying ctDNA in 4 out of 6 cases with normal PSA levels and low TF (**Fig. 7F, insert**).

We found discrepancies between the threshold model and ML models, with the ML models being unanimously accurate, while the threshold model was unanimously inaccurate (for example, 533H and 487B). There might be critical features for ctDNA calling that the threshold model missed. STREAMLINE revealed that ML models using NB, RF, XGB, and CGB all identified chrX as the first or second most important feature for binary classification (**Fig. 7G**). Because none of the samples had a seemingly significant number of top 1,000 bins on chrX, our threshold-based calling was insensitive to chrX. Therefore, we independently evaluated chrX HCBs in calling tumor cfDNA (**Fig. 7H, left**). Very few of the top 1,000 HCBs cfDNA were on chrX from control, age-matched individuals’ or buffy coat DNA from PCa patients while a much higher number of HCBs were mapped to chrX in cfDNA from PCa patients. Indeed, HCBs on chrX were statistically different between PCa cfDNA and either matched normal DNA (p = 0.013, Wilcoxon signed-rank test) or control individuals’ cfDNA (p = 0.0044, two-sided Wilcoxon rank-sum test). HCBs on chr3 were the second most influential determinant, which also showed statistically significant enrichment in PCa patients’ cfDNA (**Fig. 7H, right**).

Given that the androgen receptor gene (AR) is located on chrX and contributes to therapy-resistance in advanced PCa^53–55^, we examined AR amplification (AR-amp) through shallow WGS. AR-amp was notable in several patients with therapy-resistant tumors and very high PSA, with dozens of top 1,000 bins within the amplified regions (**Fig. 7I**). In other patients, AR-amp was not evident, and HCBs were fewer and scattered throughout chrX. These PCa cases were therapy-sensitive at the time of the blood draw but eventually progressed to resistance and life-threatening disease. HCBs on chrX, as assessed by GAPF-seq, could be an initial sign of chrX-wide rearrangement that would eventually lead to AR-amp, which could be consistent with DNA palindrome formation as an initial step of gene amplification.

## Discussion

In this study, we tested the feasibility of our genome-wide assay, named GAPF-seq, which probes DNA palindromes, for liquid biopsy-based cancer detection. DNA palindromes are mechanistic components of BFB cycles, which were shown to be a major process underlying genome instability and genomic amplification across tumor types.^36^ Consequently, targeting DNA palindromes for cancer detection seemed a rational strategy to test. GAPF is designed to enrich DNA palindromes and eliminate non-palindromic DNA. This feature would enable GAPF-seq to detect cancer-specific DNA with high sensitivity in plasma cfDNA even with low tumor fraction (TF). To validate this concept, we conducted experiments using samples with *in vitro* serial dilution of tumor DNA mixed with normal DNA as well as real-world cancer patients’ cfDNA with various TFs.

If developed as a cancer detection test, GAPF-seq would be uniquely positioned for liquid biopsy targeting SVs. While WGS and mate-pair analyses of tumor DNA can successfully identify fold-back inversions,^31,56^ it requires substantial coverage (>30x). In cfDNA, DNA representing fold-back inversions is diluted, which would demand ultra-deep coverage for accurate detection. Ultra-deep WGS is cost-prohibitive and requires significant time and effort for analyses. Targeted sequencing for gene mutation detection in cfDNA would offer a higher sensitivity for cancer detection, as exemplified by detecting tumor-informed mutations.^57,58^ However, many amplified oncogenes lack mutations. Gene amplification is not limited to treatment-naïve primary tumors for activating oncogenes; it also occurs later during disease progression to confer therapy resistance, information that is absent in primary tumors. Therefore, mutation detection tests may not be feasible for the early detection of gene amplification conferring therapy resistance. Early detection of resistance would allow physicians to consider treatment options when resistant tumors are still small. Thus, frequent monitoring through liquid biopsy and timely detection of gene amplification is important. GAPF-seq with simple cfDNA processing, low coverage NGS, and streamlined data analysis would emerge as a practical approach for frequent monitoring.

The proof of concept was tested in this study for the androgen receptor gene amplification (AR-amp) in lethal prostate cancer. AR-amp is very often associated with advanced PCa.^55^ AR signaling is a major driver of PCa progression. Contemporary AR signaling inhibitors such as enzalutamide, apalutamide, abiraterone have significantly improved the overall survival of patients in recent years.^59,60^ Despite these advancements, resistance eventually develops, leading to metastasis and lethal disease. The only clinical indication of resistance is a rising PSA. Given accumulating data supporting the benefits of early initiation of chemotherapy, a test that indicates resistance to AR suppression early would provide an opportunity to alter the therapeutic approach, potentially leading to improved survival.^53^ We found that genomic regions enriched by GAPF-seq on X chromosomes (chrX) were predominantly seen in PCa plasma cfDNA (**Fig. 7H**). Resistance cases were characterized by high TF in cfDNA and AR-amp (**Fig. 7I**). These results suggest that palindrome formation and BFB cycles underlie AR-amp. In cases that did not show AR-amp, high coverage bins (HCBs) were scattered throughout chrX. Such bins may be early signs of BFB cycles and chromosome-wide rearrangements that could eventually lead to AR-amp. If this hypothesis holds true, these patients are likely to develop AR-amp over time. If not, HCBs on chrX could be a simple indicator of cancer DNA. A comprehensive study involving longitudinally collected early and late-stage samples would distinguish these two possibilities.

We utilized chromosomal distributions of high coverage bins (HCBs) as outputs from GAPF-seq to test their efficacy in distinguishing breast tumor DNA samples from normal DNA. With our threshold model, the output performed well in most cases. However, several breast tumor (HR+) DNA and PCa cfDNA were inconsistently classified. It was noted that all of these cases exhibited low CNA levels or TF measured by ichorCNA. This suggests the possibility that tumor-derived DNA was very rare in these samples. This is most likely the case for PCa cfDNA because disease burden, estimated by PSA, was low for the inconsistently classified or GAPF-negative cases. Another potential explanation is that GAPF-seq profiles are not a robust measure of tumor DNA when chromosomal rearrangements are rare. In this regard, all three GAPF-negative cases were from HR+ tumors, which are known to be driven by hormone-dependent signaling.^61^ Understanding the discriminative power of GAPF-seq to differentiate tumor DNA from normal DNA for each tumor tissue of origin would thus be necessary.

This approach to analyzing the chromosomal distribution of the top-ranked HCBs is a zero-sum game: an accumulation of HCBs on one chromosome reduces the number of HCBs available to be distributed among the other chromosomes, which can pose challenges when interpreting autoML feature importance results. When tumor-specific DNA palindromes lead to the enrichment of HCBs on a chromosome relative to the control, it necessitates a decrease in HCBs on one or several other chromosomes. So, increases in the number of HCBs on a chromosome are likely biologically relevant (due to DNA palindromes), whereas decreases simply represent the natural result of a strictly competitive analysis rather than some decrease in palindromic content. Therefore, probabilistic algorithms such as Naïve-Bayes may be better suited for GAPF-seq over ensemble learning algorithms as Random Forest, Category Gradient Boosting, and Extreme Gradient Boosting, because enrichment of HCBs on a chromosome should have a high probability of being classified as a tumor (**Fig. 5C and 7D**).

It was somewhat unexpected that, for the breast tumor and normal pairs, the relatively straightforward threshold approach yielded superior binary classification results with respect to AUC over many ML models (**Fig. 5C**). In contrast to the ML models, which exclusively utilize HCBs as input data, the simple threshold models have the advantage of incorporating domain knowledge. This includes recognizing that (1) a burden count above some threshold may be informative and (2) there is an informative higher-level feature abstraction (i.e., one or more chromosomes have a burden above the target threshold). For ML algorithms to train a model similar to the threshold models, they would need to first learn the abstract concept that decides “IF” any of the HCB features in the dataset have a value over a target threshold. The challenge of this abstraction is reflected in long-standing computer science benchmark problems such as the “majority-on” or “even-parity” problems where simultaneous knowledge of multiple features values is required to solve the problem (i.e., the majority of features in a set above some threshold, or the sum of feature values in a set is even, respectively).^62,63^ This ML challenge can be addressed in future work via feature engineering; applying domain knowledge to construct new features that capture information across all chromosomes (e.g. construct a binary feature identifying if any chromosome in the dataset has HCBs above a given threshold). This would be expected to improve ML binary classification. Beyond feature engineering, additional opportunities to improve the predictive performance of GAPF-seq with ML include (1) utilizing more sophisticated ML algorithms, (2) conducting a more extensive optimization of algorithm hyperparameters, and (3) combining base models trained by different ML algorithms into a predictive ensemble.

On the other hand, the decision to employ chrX-based tumor/normal differentiation for PCa cfDNA/normal DNA was guided by insight from ML. Due to the scarcity of HCBs on chrX, none of the cfDNA samples reached the threshold (**Fig. 7H**). Nevertheless, the assessment of feature importance unveiled that ML algorithms strongly relied on HCBs on chrX for binary classification (**Fig. 7G**). Consequently, ML highlighted a feature that human observers (i.e., the authors) had not considered. Feature importance stands out as a widely used technique to elucidate the behavior of ML models, and its significance is particularly pronounced in the medical field.^64^ The importance of DNA palindromes on chrX is easily interpretable, given that AR-amp is a primary driver of resistance to therapy.^53–55^ These results suggest that GAPF-seq, when employing a threshold model, has limitations, and the full potential of GAPF-seq for calling tumor DNA has not been exhausted. Increasing the number of samples in future studies will enable us to enhance the analysis pipeline. Additionally, elevating the resolution of HCB positioning in 1,293 cytobands genome-wide may offer a more comprehensive assessment. For example, cfDNA fragmentation patterns were examined in 504 windows of 5 Mb genome-wide.^23,24^ DNA methylation profiling used over 1 million CpG sites in the genome, the methylation patterns of which best predict the tissue of origin and differentiate tumor DNA from normal DNA.^65^ The substantial computational power required for such analyses can be efficiently managed through the application of ML. Feature importance would then be instrumental in identifying hotspots of DNA palindromes at the cytogenetic band level.

### Limitations of the study

- We integrated cross-validation to reduce the overfitting issues of ML. Future testing with independent sample sets is necessary to support the results presented here.
- While GAPF-seq has shown efficient differentiation between tumor DNA and normal DNA for breast tumors and prostate cancer patients’ plasma, its applicability to other tumor types remains to be tested.

## Methods

### Research Samples

For mimicking DNA with low tumor fraction, we utilized the Colo320DM colorectal cancer cell line, which harbors known palindromic junctions^28^, and IMR90 normal fibroblasts. Breast cancer patients’ samples were obtained from the BioBank at Cedars-Sinai Medical Center, which stores matched frozen tumor/buffy coat/plasma samples. The project was approved by the Institutional Review Board (Pro0005127, DNA palindrome detection by liquid biopsy from breast cancer patients; 00042197 Translational Oncology Program Blood Specimen Repository). Human control plasma from single donors was procured from Innovative Research.

### DNA extraction

High Molecular Weight DNA was extracted from cell lines and tissues as described previously.^41^ Briefly, samples were incubated in 400 μL of lysis buffer (100mM NaCl, 10mM Tris, 25mM EDTA and 0.5% SDS) and 20 μL of Proteinase K (20 mg/mL) at 37 °C for 16 hours, followed by phenol/chloroform extraction and ethanol precipitation. Gene Jet Whole Blood DNA extraction kit (Thermo Scientific) was used to extract DNA from buffy coat. For plasma cfDNA extraction, a modified phenol-chloroform protocol was used. Frozen aliquots of 500 μL of plasma were thawed and mixed with 2.5 mL of cell lysis buffer and 200 μL proteinase K (20 mg/mL), followed by an incubation at 37°C for at least 16 hours. A phenol-chloroform-isoamyl alcohol mixture (Sigma-Aldrich) was added and mixed. The mixture was centrifuged at 10,000 g for 15 min and the aqueous portion was subject to a second phenol-chloroform treatment. A chloroform-isoamyl alcohol mixture ((Sigma-Aldrich) was then added and mixed, followed by centrifugation at 10,000 g for 15 min. To the aqueous portion, 0.05 mg of glycogen (Roche) and sodium acetate to a final concentration of 0.3 M were added. Then, 2:1 absolute ethanol was added and kept at −20°C overnight. The mixture was centrifuged at 14,000 g for 30 min at 4°C and the pellet was washed with 70% ethanol before being resuspended in low TE buffer (Thermo Scientific). cfDNA concentration was measured using dsDNA HS Assay Kit and a Qubit 3.0 fluorometer (Life Technologies). DNA fragment analysis was performed using Agilent 2100 Bioanalyzer system (Agilent).

### Genome-wide Analysis of Palindrome Formation

For the analysis of small amounts of DNA, we devised a modified protocol of the Genome-wide Analysis of Palindrome Formation (GAPF)^38^. Briefly, either 100 ng or 30 ng of input genomic DNA was digested with either KpnI or SbfI, both of which are relatively infrequent cutting enzymes, at 37°C for 16 hr. The restriction enzymes were heat-inactivated at 65°C for 20 min and the reaction products were mixed. This mixture was boiled for 7 min in 100 mM NaCl and 50% formamide, then immediately quenched in ice water for 5 min. Subsequently, DNA was digested with 100 U/μg of S1 nuclease (Roche) at 37°C for 1 hour and was processed with the ChargeSwitch PCR Clean-up Kit (Invitrogen) or Monarch PCR and DNA Cleanup Kit (New England Biolabs).

The purified DNA was fragmented using the Covaris M220 and the libraries were constructed using the NEBNext Ultra II DNA Library Prep Kit (New England Biolabs) for next generation sequencing. Alternatively, GAPF libraries were constructed with the NEBNext Ultra II FS DNA Library Prep Kit (New England Biolabs), which employs fragmentase for DNA fragmentation. Whole-genome sequencing libraries were constructed using 40 ng of genomic DNA.

### Whole Genome Sequencing

All samples were sequenced on the Illumina HiSeq System by Novogene Co., Ltd. to generate 150 bp paired-end sequencing reads, achieving approximately 7.5x coverage. The reads underwent adaptor removal using Trim Galore (v0.6.1) and Cut Adapt (v2.3) (*trim galore options: --length 55*) and aligned to the UCSC hg38 human reference genome (hg38.fa.gz, 2014-01-15 21:14) using the DNA alignment software Bowtie 2 (v2.3.5) (*options: -x -U -S or -x -1 -2 -S*). The aligned reads were converted to binary format (*samtools view options: -bS*), reordered (*samtools sort options: -o*), and sorted (*sort options: -k1,1 -k2,2n -o*) using Samtools (v1.9) and counted in genome-wide 1 kb non-overlapping bins using Bedtools (v2.28.0) (*bedtools coverage options: -sorted -counts -a -b -g*). Coverage was visualized using the Integrative Genomics Viewer (IGV) (v2.5.0). Read counts were normalized by a per-million scaling factor to adjust for the sequencing depth.

### GAPF-seq Quality Control

To assess the quality of palindrome amplification by GAPF-seq, we examined the read coverage for known inverted repeats on chromosomes 1, 9, 17, 19, and X (**Supplementary Table 1**). These short inverted repeats are infrequently cut by the restriction enzymes and readily amplified by GAPF. Reads were counted at specific inverted repeats at the X chromosome genes DMRTC1, SSX2, SSX4, CXorf49, CXorf51A, PNMA6A, and SPANXA2 as well as the inverted repeats at HIST2H3C on chr1, FAM225A on chr9, SNORD3A on chr17, and 19p13.2. Normalized read counts for each of the regions were averaged to derive an amplification score. Because these regions are repeated within the reference genome, filtering based on mapping quality score that only returns reads that uniquely map to the reference genome should deplete the normalized read counts (*samtools view options: -b -q 40*). Thus, filtered values for each region were averaged to determine a depletion score. GAPF-seq passes quality control with an amplification score above 1.0 and a depletion score less than 0.075, which indicates the enrichment of DNA palindromes and elimination of non-palindromic or repeated sequences.

### GAPF Background Bins

Due to their repetitive structure, high GC content and epigenetic modifications, certain regions of the genome may yield false positives or obscure the identities of DNA palindromes.^38^ To mitigate this, we first applied a filter based on mapping quality score (MAPQ > 40), retaining reads that uniquely map to the reference genome (hg38). Subsequently, we excluded regions that were commonly enriched in both tumor and control samples. This curation resulted in 3,133,748 bins for further GAPF analysis, down from the original 3,209,513 non-overlapping 1 kb bins (**Supplementary Table 2**).

### Cytoband Analysis

Coordinates for cytogenetic bands in hg38 were downloaded from the UCSC genome browser (cytoBand.txt.gz, 2022-10-28 21:15). Following the removal of the background bins (**Supplementary Table 2**), reads were counted in cytobands in order to calculate the coverage for each cytoband. There were 1,293 cytobands downloaded from the UCSC genome browser. Unlocalized sequences were removed from the analysis leaving a total of 863 cytobands for subsequent analyses. For figures containing 1-kb HCBs in cytobands, the number of top 1,000 high coverage GAPF bins corresponding to each cytoband were counted.

### Tumor Fraction

DNA copy number (CN) and tumor fraction (TF) were computed from shallow whole genome sequencing data using the hidden Markov model implemented in the software package ichorCNA (v0.3.2).^66^ (*read counter options: --window 1000000 --quality 20 --chromosome “chr1, chr2, chr3, chr4, chr5, chr6, chr7, chr8, chr9, chr10, chr11, chr12, chr13, chr14, chr15, chr16, chr17, chr18, chr19, chr20, chr21, chr22, chrX, chrY”*) (*runIchorCNA options: --ploidy “c(2,3)” --normal “c(0.5,0.6,0.7,0.8,0.9)” --maxCN 3 --gcWig gc_hg38_1000kb.wig --mapWig map_hg38_1000kb.wig --centromere GRCh38.GCA_000001405.2_centromere_acen.txt --normalPanel HD_ULP_PoN_hg38_1Mb_median_normAutosome_median.rds --includeHOMD False --chrs “c(1:22, \”X\”)” --chrTrain “c(1:22)” --estimateNormal True --estimatePloidy True --estimateScPrevalence False --scStates “c()” ---txnE 0.9999 --txnStrength 10000*)

### GAPF Binary Classifier (Threshold Model)

The normalized read coverage in 1 kb non-overlapping bins following the mapping quality score filter (MAPQ score >= 40) were sorted in descending order (*sort options: -nr -k 4 -k 2n*). The top 1,000 bins according to coverage were considered for subsequent analyses (*head options: -1000*). The number of these high coverage bins (HCBs) corresponding to each chromosome were counted (totaling 1,000). A binary classifier designated samples as GAPF-positive or GAPF-negative according to the number of HCBs on a chromosome against a threshold. Using the 39 breast tumor and control samples, the sensitivity and specificity were calculated at various thresholds for HCBs on a chromosome. A receiver operating characteristic curve (ROC) was generated and the area under the curve (AUC) was calculated using the trapezoidal rule. The optimal threshold was determined using Youden’s J statistic.

### Automated Machine Learning Pipeline (STREAMLINE)

Machine learning binary classification modeling and evaluation was conducted using version 0.2.5 of the Simple Transparent End-To-End Automated Machine Learning Pipeline (STREAMLINE),^48^ applying mostly default pipeline run parameters (exceptions identified below). Input data were the number of top 1,000 HCBs on chromosomes 1-22 and X (23 features) unless otherwise stated. Three-fold cross-validation (CV) was used in the predictive modeling using the following classification algorithms: Naïve-Bayes (NB), Random Forest (RF), Extreme Gradient Boosting (XGB),^67^ Category Gradient Boosting (CGB),^68^ or Extended Supervised Tracking and Classifying System (ExSTraCS).^69^ Within STREAMLINE, all algorithms with hyperparameters (excluding NB) underwent an Optuna-driven hyperparameter sweep^70^ with 200 target trials and a timeout of 15 minutes across a broad range of hyperparameter options. In addition to evaluating all models across 16 classification metrics, STREAMLINE also calculates model feature importance estimates uniformly for each algorithm and model using permutation feature importance estimation. STREAMLINE model training was repeated for each analysis using random seeds of 22, 32, or 42 unless otherwise stated.

## Supporting information

Supp table 1 GAPF QC regions

Supp table 2 GAPF bkgd regions

Supp table 3 raw data breast tumor

Supp table 4 raw data breast duplicates

Supp table 5 raw data breast tumor 300 HCBs

Supp table 6 raw data breast tumor 10k HCBs

Supp table 7 raw data breast tumor diluted

Supp table 8 raw data prostate cfDNA

Supp table 9 raw data control cfDNA

## Acknowledgement

This work is supported by the National Cancer Institute (2 <u>R01 CA149385</u>), Department of Defense (W81XWH-18-1-0058), and Cedars-Sinai Medical Center (to H.T.); the Margie and Robert E. Petersen Foundation (to A.E.G.).

## Author Contributions

M.M.M., A.E.G. and H.T. conceived and designed the experiments. M.M.M., F.I. and L.M. generated genomic data. M.M.M., F.I. and H.T. designed the data analysis workflow. M.M.M. and S.K. performed statistical analyses. R.U. designed the ML pipeline software, interpreted the analyses, and revised the manuscript. Z.C. and E.M.P. provided prostate cancer samples and interpreted the data. M.M.M. and H.T. wrote the manuscript. D.D., A.E.G. and H.T. supervised the project.

