## Supplementary Figures for "A Practical Approach for Targeting Structural Variants Genome-wide in Plasma Cell-free DNA"

### Supp Figure 1, Murata et al.

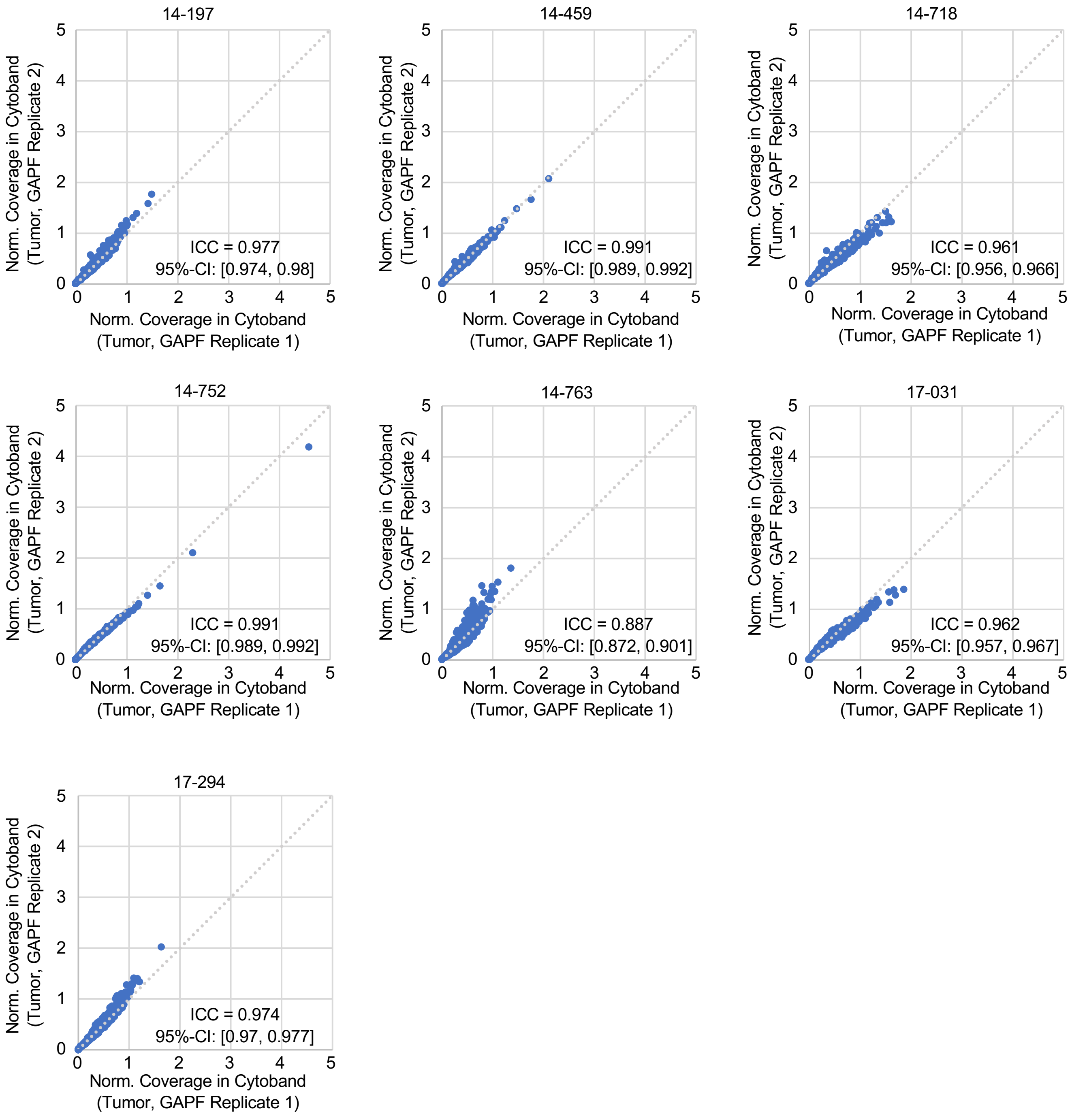


**Supp Fig. 1: GAPF-seq is Reproducible**. The GAPF read coverage in cytobands is normalized by the size of the cytoband (in bp) for the seven duplicates individually. The normalized read coverage in each cytoband for the first replicate is plotted against the duplicate. The overall intraclass correlation coefficient is 0.964 (95% CI: 0.962-0.966).

### Supp Figure 2, Murata et al.

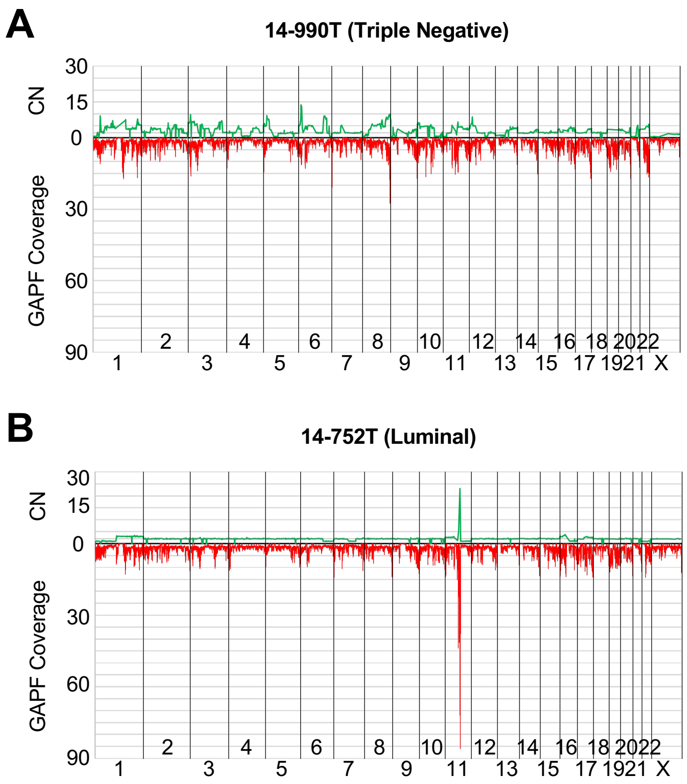


**Supp Fig. 2: DNA Palindromes Correlate to CNA**. The copy number (CN) is plotted against the GAPF read coverage for A) a triple negative breast cancer (TNBC) tumor 14-990 and B) a luminal (Lum) breast tumor 14-752.

Supp Figure 3, Murata et al.

A

B

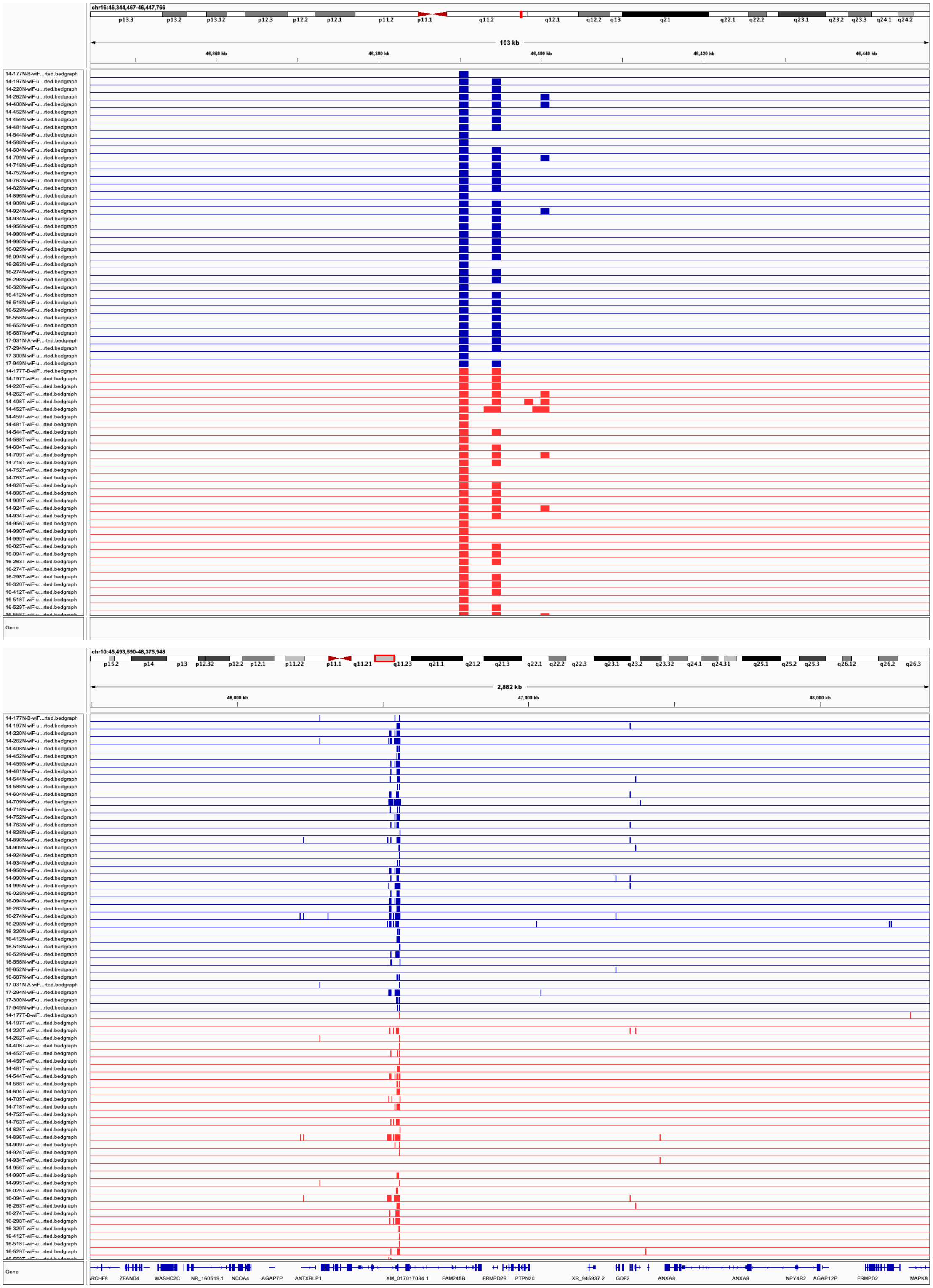

**Supp Fig. 3: Bins Commonly Enriched in Both Tumor and Normal Samples Are Removed from Analyses**. Certain regions of the genome may be enriched by GAPF-seq, but do not represent *de novo* palindromes. Instead, they may harbor GC-rich sequences or repetitive elements and can be removed from subsequent analyses such as these regions from  A) chr16 and B) chr10.

### Supp Figure 4, Murata et al.

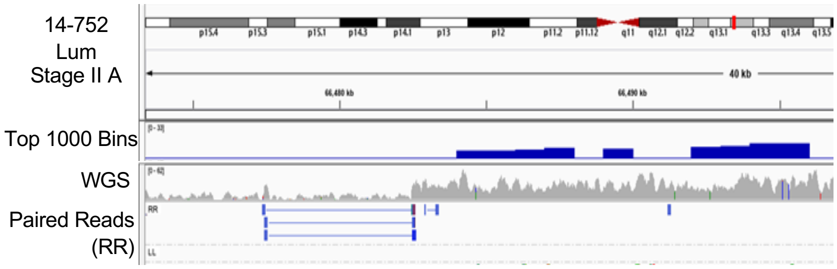


**Supp Fig. 4: Paired Read Orientations Demarcate Palindrome Junctions**. The top 1,000 bins, shallow whole genome sequencing (WGS) data, and paired read orientations visualized in Integrative Genomics Viewer (IGV). "RR" oriented read pairs are found at the borders of copy number alterations and top 1,000 HCBs. The mechanism of how palindromes create "RR" read pairs is shown in Fig. 1D.

Supp Figure 5, Murata et al.

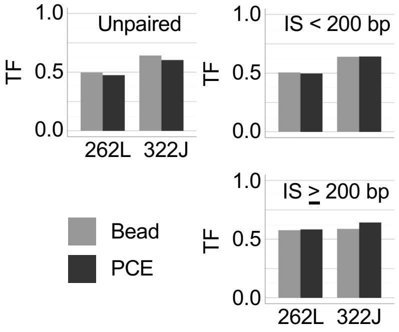


**Supp Fig. 5: Long-fragment cfDNA Harbors ctDNA Based on Insert Size**. Paired-end alignment of sequencing reads can differentiate cfDNA populations by fragment length based on the insert size. Long-fragment cfDNA extracted by phenol-chloroform (PCE) would likely result in read pairs with insert sizes greater than 200 bp whereas short fragment cfDNA extracted by beads (Bead) would likely result in insert sizes less than 200 bp. In two plasma samples, the tumor fraction (TF) was equally represented after filtering read pairs by insert size.
